# New Soluble Angiopoietin Analog of C4BP-ANG1 Prevents Pathological Vascular Leakage

**DOI:** 10.1101/2020.03.30.016147

**Authors:** Pan Liu, Michael Ryczko, Xinfang Xie, Aftab Taiyab, Heather Sheardown, Susan E. Quaggin, Jing Jin

**Affiliations:** Feinberg Cardiovascular and Renal Research Institute, Department of Medicine/Nephrology, Feinberg School of Medicine, Northwestern University, Chicago, IL 60611; Mannin Research, Inc., Toronto, Ontario, Canada, M5G 0B7; Department of Nephrology, The First Affiliated Hospital of Medical College, Xi’an Jiaotong University, Xi’an, China; Department of Chemical Engineering, McMaster University, 1280 Main St W, Hamilton, ON L8S 4L8, Canada

**Keywords:** Angiopoetin 1, Angiopoietin-Tie2 pathway, Vascular leak, chimeric protein

## Abstract

Vascular leak is a key driver of organ injury in diseases such as Acute Respiratory Distress Syndrome caused by viruses, including COVID-19. Strategies that reduce enhanced permeability and vascular inflammation are promising therapeutic targets. Activation of the Angiopoietin-1 (Angpt1)-Tie2 tyrosine kinase signaling pathway is an important regulator of vascular quiescence. Here we describe the design and construction of a new soluble ANGPT1 mimetic that is a potent activator of endothelial Tie2 in vitro and in vivo. Using a chimeric fusion strategy, we replaced the extracellular matrix (ECM) binding and oligomerization domain of ANGPT1 with a heptameric scaffold derived from the C-terminus of serum complement protein C4-binding protein α (C4BP). We refer to this new fusion protein biologic as C4BP-ANG1, which forms a stable heptamer and induces TIE2 phosphorylation in cultured cells, and in the lung following *i.v.* injection of mice. Injection of C4BP-ANG1 ameliorates VEGF- and lipopolysaccharide-induced vascular leakage, in keeping with the known functions of Angpt1-Tie2 in maintaining quiescent vascular stability, and therefore is a promising candidate treatment for inflammatory endothelial dysfunction.

## Introduction

Since the first clinical use of a VEGF antagonist for cancer and vascular eye disease more than a decade ago(1), there has been rapid development of new drugs targeting the VEGF-VEGFR2 and Angpt-TIE2 pathways(2). In the Angpt-Tie2 pathway the major ligands are ANGPT1, which is a strong Tie2 agonist, and ANGPT2, which acts as a context-dependent antagonist of Tie2 (3, 4). One of the key distinctions between these two ligands is that ANGPT1 self-clusters into higher order oligomers, and this is critical to induce Tie2 activation (5, 6). ANGPT1 is secreted by a variety of perivascular cell types to act on its cognate receptor Tie2 on the cell surface of endothelium, where it regulates junction stability to protect against vascular leakage. Localized or transient increases in blood vessel permeability occur during inflammation, ischemia, and tumor-related neoangiogenesis (7–9). Because the reduction of vascular leak has been linked to better patient outcomes in sepsis, ARDS and other conditions, therapeutic targeting of the Angpt-Tie2 pathway has been attempted by many groups (2, 10–15).

Currently a number of investigational drugs are being tested that are aimed at activating Tie2 activity (2). These include the recombinant proteins BowAng1 (16) and COMP-ANG1 (17), where the Tie2-binding fibrinogen-like domain (FLD) of ANGPT1 is fused to either the Fc of IgG1 (BowAng1) or the coiled-coil domain (CCD) of rat cartilage oligomeric matrix protein (COMP), respectively. Small molecules that inhibit the activity of TIE2 phosphatase VE-PTP are able to potently activate Tie2 in a ligand independent manner (18, 19). In addition, therapeutic antibodies have been developed against ANGPT2 that antagonizes Tie2 in the inflamed vasculature. These include ANGPT2 neutralization antibodies such as trebananib, MEDI3617, LY3127804, nesvacumab and REGN910-3, as well as bispecific ANGPT2-VEGF-targeted antibodies vanucizumab/RG7221 and RG7216 (2, 20–22). More recently, ANGPT2-targeted antibody ABTAA has been shown to trigger ANGPT2 clustering, converting ANGPT2 into an ‘ANGPT1-like’ agonist of TIE2 (23). To date, ANG-TIE-targeted drugs have been tested in patients with cancer as well as in patients with retinal vascular diseases such as wet age-related macular degeneration (wAMD) and diabetic macular oedema (DMO) (2). In all of these clinical settings, vascular leak is believed to play a pathogenic role.

Mode of delivery is an important consideration of Angpt-Tie-targeted therapies. In addition to binding Tie2 at the cell-cell contacts and stabilizing endothelial junctions, ANGPT1 has an affinity to bind to a variety of extracellular matrices (ECM) (24), responsible for anchoring endothelial Tie2 to cell-ECM attachments. As the action(s) of endogenous ANGPT1 is largely restricted to mediating pericyte-endothelium communication, in order to create a systemic therapy to activate Tie2, we needed to consider how to overcome the ‘stickiness’ or affinity of ANGPT1 to rapidly bind ECM and become immobile. To overcome the issue of ECM binding, we replaced the coiled-coil domain and linker sequence of ANGPT1 with a 57 amino acid segment derived from serum complement C4BP that naturally folds into a heptavalent scaffold, which allows the recombinant protein to circulate freely while maintaining potent Tie2 activating ability.

## Results

### Construction of C4BP-ANG1, a recombinant ANG1 mimetic that activates TIE2

In contrast to native ANGPT1 comprised of an N-terminus supercluster domain (SCD) and coiled-coil domain (CCOD) followed by a Tie2-binding fibrinogen-like domain (FLD) (**Fig1A**), two recombinant fusions between ANGPT1 FLD (amino acid 281-498) and the C-terminus domain of C4BPα (amino acid 541-597) were constructed as C4BP-ANG1 and ANG1-C4BP in their N- to C-terminus orders and produced using HEK293 and CHO cell production systems (Methods). It is expected that the C4BP segment forms a seven-unit stalk further secured by intermolecular disulfide bridges (**Fig1B**, top), which cluster seven FLDs of ANGPT1 in a stable molecular complex (**Fig1B**, bottom). In keeping, both C4BP-ANG1 and ANG1-C4BP appeared as high order oligomers of ~250 kDa in size on non-reducing SDS-PAGE, in contrast to their reduced monomeric counterparts of ~35 kDa (**Fig1C**). Electron microscopy further confirmed the formation of C4BP-ANG1 heptamers (**Fig1D**). Because storage stability of ANGPT1-based biologics tends to be problematic (25), we expressed and tested several variants of the fusion constructions. The results showed regardless of the N-to-C domain order and 6xHis tag location all C4BP-ANG1 variants formed stable complexes (supplementary Figure S1), and could be safely stored in physiological buffer conditions with no formation of protein aggregates. In addition, we specifically tested ANG1-C4BP after freeze-thaw cycles. There was no quantity or quality decline of the protein complex as determined by UPLC-SEC analysis (supplementary Figure S2).

**Figure 1.**
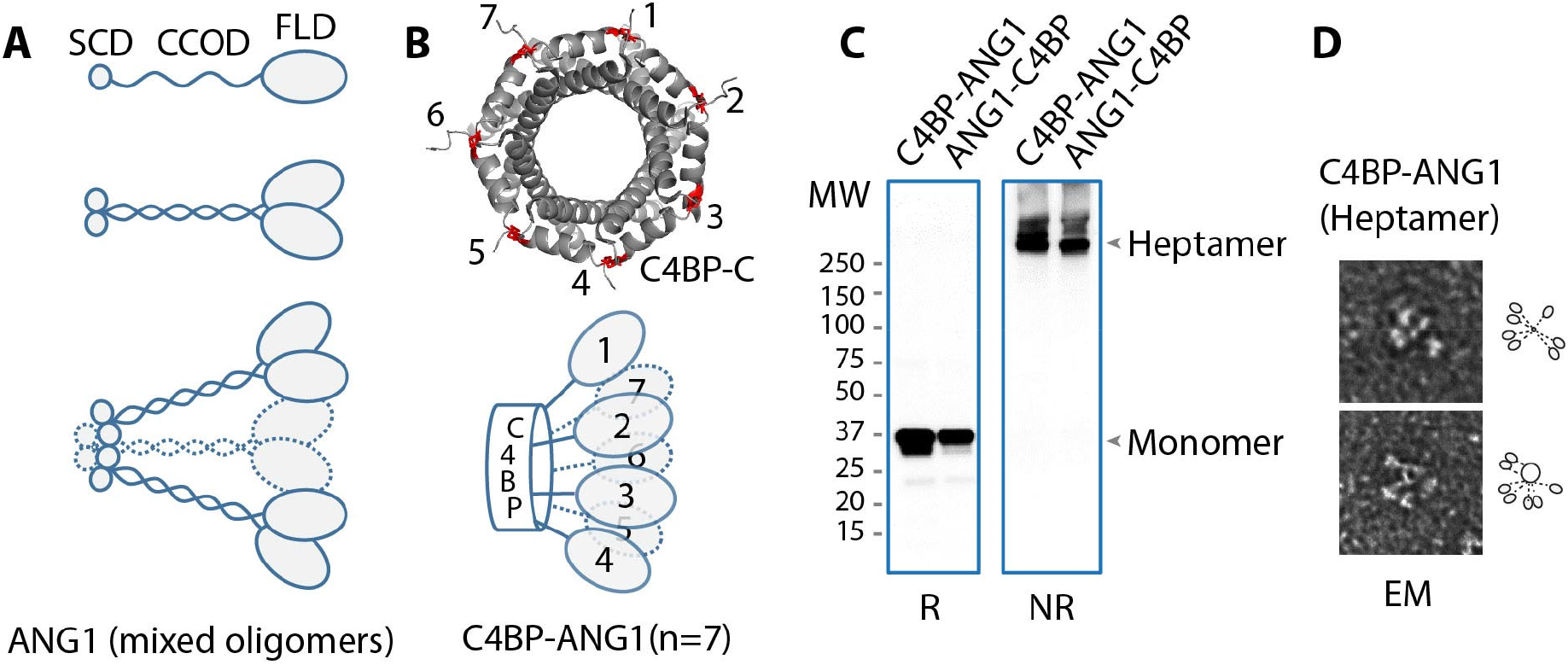
Construction of heptameric C4BP-ANG1. **A.** Native ANG1 is comprised of, from an N- to C- terminus order, a supercluster domain (SCD), a coiled-coil domain (CCOD), and a fibrinogen-like domain that binds Tie2 (top). The CCOD mediates CCOD-CCOD interactions between ANG1 molecules (middle), and through its linker segment with FLD, also binds the ECM. The SCD further clusters ANG1 into higher degree complexes (bottom). **B**. The C-terminus of C4BP naturally folds into a “barrel” structure through inter-linking disulfide bridges (red) between neighboring subunits. A total of seven (or eight) of these subunits complete the barrel structure (top) that, in C4BP-ANG1 or ANG1-C4BP, displays seven FLD in an arrangement reminiscent of that of native ANG1 (bottom, compared to A). **C**. C4BP-ANG1 and ANG1-C4BP was expressed through transfection of the encoding plasmid to HEK293 cells and purified from culture media. Both proteins were detected as monomers on SDS-PAGE under reducing condition and heptamers under non-reducing (NR) condition. **D**. Electron micrograph (EM) images showed clustered C4BP-ANG1.

### C4BP-ANG1 activates TIE2 in cultured cells

To determine direct binding between C4BP-ANG1 and Tie2, in a series of co-immunoprecipitation assays we showed binding of C4BP-ANG1 and ANG1-C4BP to Tie2-Fc (**Fig2A**). As expected, by adding purified C4BP-ANG1 or ANG1-C4BP to culture medium of human umbilical vein cells (HUVECs) we observed activation of Tie2 as measured by phospho-Akt (**Fig2B**). Using a human embryonic kidney (HEK293) cell system that stably expressed full-length Tie2, we showed both C4BP-ANG1 and ANG-C4BP induced Tie2 phosphorylation in a dose-dependent manner (**Fig2C**). Furthermore, immunofluorescence studies showed a dynamic redistribution pattern of Tie2 on the surface of HUVECs following C4BP-ANG1 stimulation, resembling the effects of natural ANGPT1 (26). C4BP-ANG1 caused Tie2 to coalesce into small puncta that also migrated towards the cell periphery (**Fig2D**), further suggesting adequate signaling through Tie2 is achieved. As expected, confluent HUVECs in culture formed continuous cell-cell junctions (**Fig2E**), whereas treatment of the cells with either LPS or VEGF caused disruption of the junctions in some areas. Meanwhile, when C4BP-ANG1 was present, junctional integrity was maintained in spite of LPS and VEGF treatments (**Fig2E**), and this is consistent with the function of ANGPT1 in maintaining endothelial quiescence and stability.

**Figure 2.**
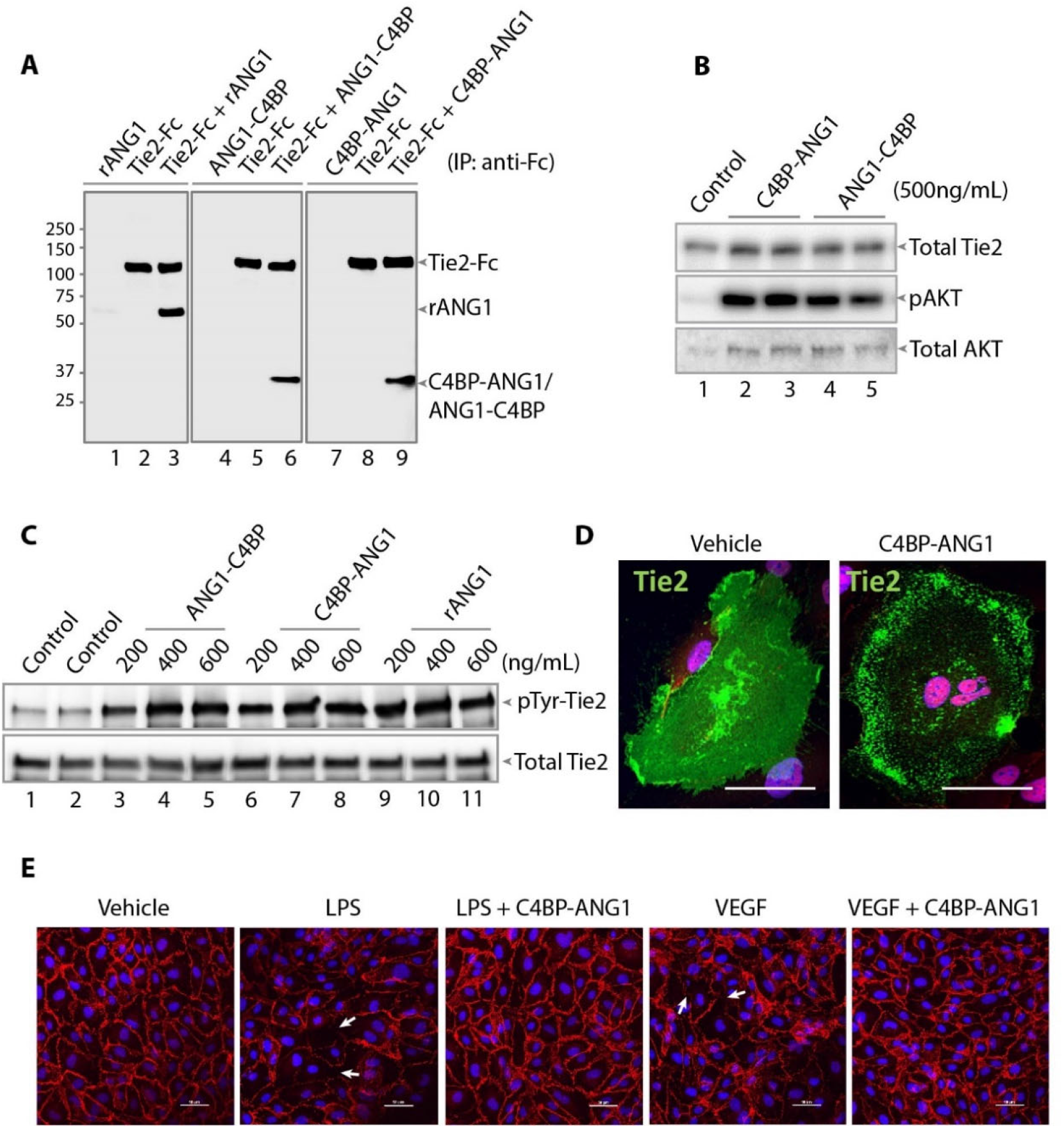
*In vitro* activity of C4BP-ANG1 towards Tie2. **A.** Using the ectodomain of Tie2 in the form of an Fc fusion (Tie2-Fc), direct interactions between Tie2 and recombinant ANG1 (rANG1) of native ANG1 sequence, C4BP-ANG1 or ANG1-C4BP were tested in a co-immunoprecipitation assay. Following anti-Fc immunoprecipitation, the presence of the ANG1 variants with Tie2-Fc were detected by anti-His tag blotting. **B**. Both C4BP-ANG1 and ANG1-C4BP in alternative N-to-C domain arrangements could activate HUVEC cells as measured by AKT phosphorylation (pAKT). **C**. Using a HEK293 cell line that had been stably transfected with FLAG-Tie2 (full length), C4BP-ANG1 treatments of the cells elevated phospho-tyrosine (pTyr) levels as measured by immunoprecipitation with anti-FLAG antibody followed by immunoblotting with anti-pTyr antibody. **D**. HUVECs were transfected with FLAG-Tie2 (full length) and subjected to vehicle control or C4BP-ANG1 treatment. The subcellular distribution patterns of Tie2 were shown by anti-FLAG immunofluorescence staining (a representative single cell image is shown in each group). **E**. Confluent HUVECs were treated with LPS, LPS together with C4BP-ANG1, VEGF, or VEGF together with C4BP-ANG1 for 30 min. Following fixation, the cells were stained with VE-Cadherin (in red) that marks cell junctions. While LPS and VEGF causes partial disruption of junctions (pointed by arrows), co-treatment of the cells with C4BP-ANG1 maintained the integrity of the junctions. Scale bar: 50 μm.

### *In vivo* pharmacoefficacy of C4BP-ANG1

With as little as 0.2 μg/g (in body weight) of purified C4BP-ANG1 was injected intravenously in mice, elevated levels of phospho-Tie2 were detected in the lung (**Fig3A**). A strong phospho-Tie2 signal persisted for at least 6 hours following a bolus injection of C4BP-ANG1, and residual Tie2 activities remained detectable 16 hours later (**Fig3B** and **3C**). Intraperitoneal injection of young mice also resulted in Tie2 activation in the lung (**Fig3D**). Since vascular therapies have broad indications for ocular diseases (27), we also tested pharmacokinetics of the compound following intraocular injection in rabbits. Following a single intravitreal injection of C4BP-ANG1, the presence of the drug was detected to last for at least 5 days in the aqueous (**Fig3E**), and 7 days in the vitreous (**Fig3F**), demonstrating excellent *in vivo* stability of the biologic.

**Figure 3.**
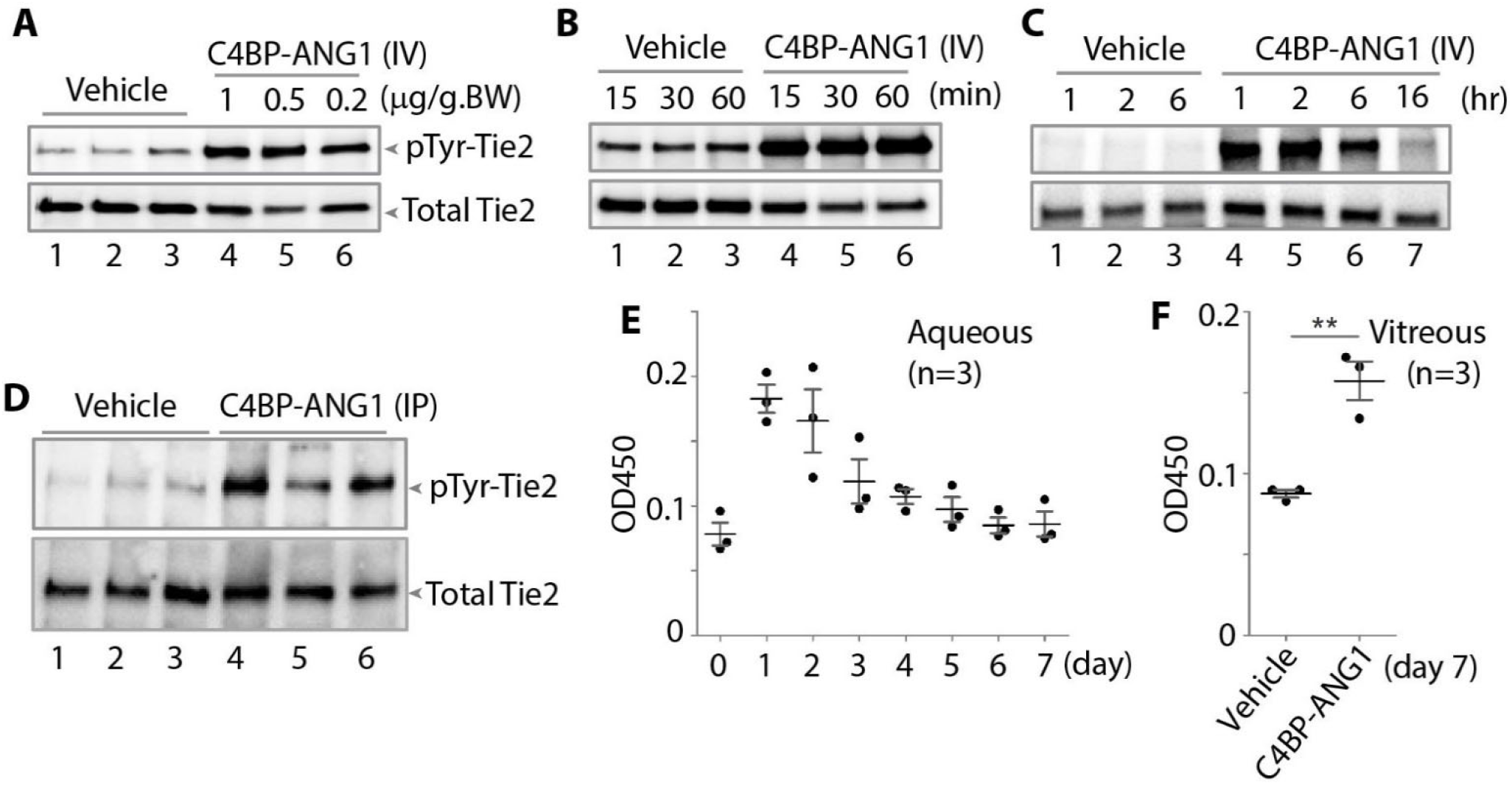
*In vivo* injection studies of C4BP‐ANG1. Mice (A‐D) and rabbits (E‐F) were injected with C4BP‐ ANG1 and *in vivo* activities were measured either by phosphorylation of endogenous Tie2 in the lung (A-D) or by the levels of C4BP‐ANG1 in the eye (E‐F). **A.** Mice were *i.v.* injected with either vehicle or C4BP‐ ANG1 of different doses based on body weight (BW) and lung tissues were harvested 30 min after. Following anti‐Tie2 immunoprecipitation, phospho‐Tie2 levels were measured by immunoblotting with anti‐pTyr antibody. **B** and **C.** time course studies of phospho‐Tie2 in response to C4BP‐ANG1 at 0.5 μg/g.BW. **D.** P5 neonatal mice were injected with C4BP‐ANG1 (0.5 μg/g.BW.), or vehicle control, through intraperitoneal (IP) administration and tissues were harvested four hours later for measuring phospho‐ Tie2 levels. **E.** Three rabbits were each subjected to a single dose of intravitreal injection of C4BP‐ANG1 and aqueous humor was collected daily (preinjection sample: day 0) for seven days. The levels of C4BP‐ANG1 were measured by ELISA using anti‐His capturing antibody and anti‐ANG1 detection antibody (OD450 values). **F.** On the seventh day the animals were sacrificed and vitreous samples were collected for detecting C4BP‐ANG1 levels (asterisks: *p*<0.01).

### C4BP-ANG1 ameliorates pathological vascular leakage

Next, we performed a series of pharmacologic tests using mouse models, in which we evaluated the effects of the drugs in protecting mice from pathologic vascular leakage (**Fig4**). By using Evan’s Blue tracer, we examined the *in vivo* efficacy of C4BP-ANG1 in preventing VEGF-induced microvascular leakage in the skin. The experiments demonstrated C4BP, either administrated locally through intradermal injection (**Fig4A**), or systemically through *i.v.* injection (**Fig4B**), markedly reduced leakage at the locations of intradermal VEGF injection. Furthermore, in a chemical-induced vascular leakage model using mustard oil, the levels of Evan’s Blue leakage were also significantly lower in the C4BP-ANG1 treatment group (**Fig4C**). To model lung injury leading to ARDS in sepsis, we adapted a lipopolysaccharide-inhalation protocol to induce lung edema. The mice were divided into groups with or without being treated with C4BP-ANG1 from *i.v.* injection. As expected, mice that received two doses of C4BP-ANG1 treatments were partially protected from lipopolysaccharide-induced lung injury as measured by vascular leakage (**Fig4D**). Meanwhile, in a tolerance test with repeated doses, we injected mice daily for 14 days. There were no overt adverse effects (not shown).

**Figure 4.**
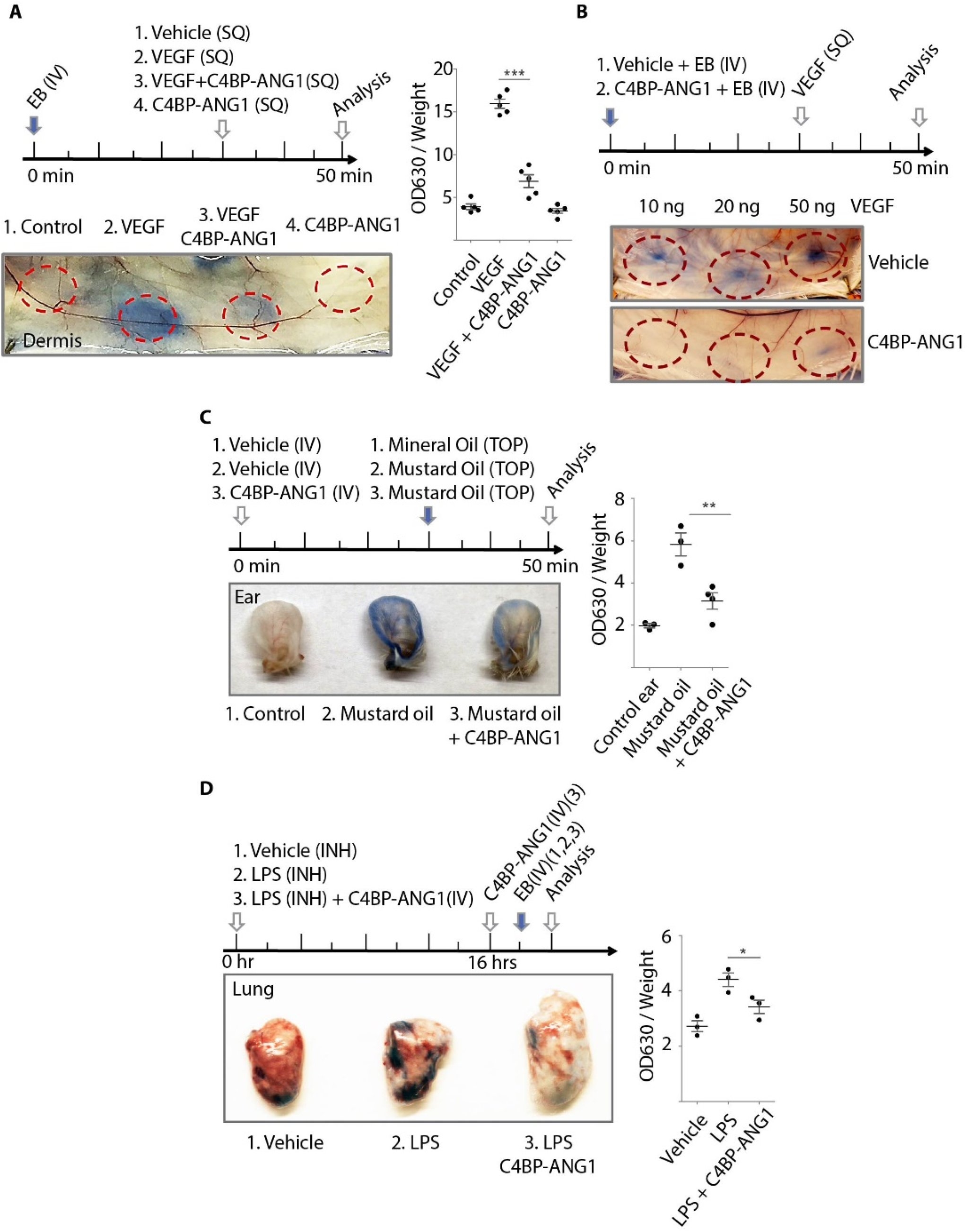
C4BP‐ANG1 injection ameliorates pathologic vascular leakage. The studies of vascular leakage were conducted using Miles assay that quantifies tissue levels of Evans Blue dye. **A**. Mice were subjected to a 30 min injection schedule as shown (top). Subcutaneous (SQ) injections of a combination of VEGF and C4BP‐ANG1 were performed and leakage of Evans Blue (EB) were visualized (bottom) and quantified as OD360 values normalized by tissue weight (right. asterisks: *p*<0.001). **B** and **C**. Instead of local injection of C4BP‐ANG1, the biologic was administrated prophylactically via *i.v.* 30 min prior to leakage induction either by VEGF (B) or by mustard oil (C. asterisks: *p*<0.01). **D**. In a mouse model of lipopolysaccharide (LPS)‐ induced lung injury, a time course of LPS inhalation (INH), C4BP‐ANG1 injection and Evans Blue injection was followed as indicated in the top panel. One hour after Evans Blue injection, the lung was harvested for measuring leakage (image and quantification. asterisk: *p*<0.05).

## Discussion

Here we sought to develop a “bio-better” analog of ANGPT1 as pharmaceutical candidate using a rational recombinant engineering design. Our overall strategy was to overcome a key therapeutic limitation of native ANGPT1, which is nonspecific sequestration by the ECM and easy to aggregate, while still to preserve a high degree of oligomerization. We chose a protein scaffold segment from C4BP, an abundant serum complement protein complex, which is characterized by seven functional polypeptide chains extending from an intertwined scaffold (28). We constructed several variants of C4BP-ANG1 fusion proteins and showed all formed stable heptamers when expressed in HEK293 and CHO cells. Intradermal, intravenous, intraperitoneal and intravitreal injections of mice and rabbits all showed long-lasting activity and protection from a variety types of inflammatory vascular leak.

In recent years, there have been major advances in drug development that specifically target the Angpt-Tie pathway. Strategies include ANGPT1 mimetics, ANGPT2 neutralizing antibodies, bi-specific, double-targeting antiangiogenic antibodies or affinity traps linking ANGPT2 with VEGF, or similarly, ANGPT2 with Tie2, and small molecule inhibitors of VEPTP or Tie2 catalytic activities, among others(2). With the exception of Tie2 kinase inhibitors, all these therapies are designed to elevate the activity and function of Tie2 to promote vascular stabilization and reduce vascular response to inflammatory signals. Theoretically, these activities should counteract VEGF-induced neovascularization and inflammatory leakage that is characteristic of many human diseases. Many clinical studies have demonstrated an association between elevated expression of circulating ANGPT2 and poor clinical outcome in diseases characterized by vascular leakage and tissue damage (2, 29). While it remains unclear if these elevated levels of ANGPT2 antagonizing Tie2 activity during inflammation are causal, agonistic ANGPT1 mimetic treatment with C4BP-ANG1 is predicted to ameliorate vascular destabilization that occurs in severe infectious conditions such as pneumonia from flu or coronavirus. Indeed, several clinical studies confirm that the ratio of circulating ANGPT1/ANGPT2 is predictive of patient outcome (30, 31).

Prior to this study, ANGPT1-mimetic therapies have been designed and tried where the N-terminus domains of the ANGPT1 analogs have been removed and replaced with less “sticky” structures. These strategies include BowAng1, which contains a dimerization Fc domain and a tandem arrangement of the FLD, achieving a multimericity of 2×2=4. A second strategy fused the FLD of ANGPT1 to the short coiled-coil domain of cartilage oligomeric matrix protein (COMP), which generated a pentameric complex with “super ligand” activity. However, while COMP-ANG1 has improved solubility over native ANG1, the coiled-coil-to-coiled-coil interactions, which tend to have broad interaction preference among different coiled-coil domains (32, 33), have the propensity to trigger protein aggregation during production (15, 25). It may also retain at the ECM after injection due to a general affinity of the coiled-coil domain towards tissue coiled-coil proteins that are abundantly found in the ECM (24, 34). We hypothesized that the selection of a scaffold with better serum-compatibility may solve multiple problems: 1) ECM-sequestration 2) degradation from systemic circulation 3) protein aggregation during production and storage and 4) possible antigenicity. Besides forming a heptameric scaffold, the selected C4BP complex used to construct C4BP-ANG1 does not carry immunogenicity of the parent protein. Hence, while C4BP-ANG1 structurally resembles native C4BP heptameric complex, it is able to function as a potent Tie2 ligand via its highly clustered FLDs. Furthermore, C4BP-ANG1 is well tolerated in mice with daily injections for two weeks. Since the fusion was constructed using all human protein sequences of C4BP and ANG1, we have not performed long-term injection studies beyond 14 days due to the concern of development of cross-species neutralizing antibodies.

In summary, we designed and constructed a new C4BP-ANG1 fusion as a new candidate ANGPT1 mimetic therapy. The fusion displays remarkable stability with high multimericity that can potently activate Tie2 and is suitable for systemic administration. Furthermore, the study validated a general strategy to use a C4BP scaffold in drug design when high degree of oligomeric states is desired.

## Materials and Methods

### Construction of C4BP‐ANG1 cDNA

ANG1‐C4BP and C4BP‐ANG1 refer to chimeric fusions between ANGPT1 C‐terminus FLD and C4BP C‐terminus segments in an N‐to‐C‐terminus order, respectively, in either direction. The C4BP C‐terminus segment is derived from human C4BPα protein (Accession No. NP_000706.1). The C‐terminus FLD segment of ANGPT1 is derived from the human sequence (Accession No. NP_001137.2; amino acid 281‐ 498). DNA sequences encoding ANG1‐C4BP and C4BP‐ANG1 were synthesized by Integrated DNA Technologies and cloned into pcDNA3 vector, a poly‐histidine tag was included on the C‐termini of the sequences.

### Recombinant protein production and purification

Production of recombinant C4BP‐ANG1 and ANG1‐C4BP was performed in HEK293 clonal expression system that was described previously (35). Briefly, the constructs in expression vector pcDNA3 were transfected to HEK293 cells, and selected in DMEM medium supplemented with 800 μg/ml of G418 until individual colonies were visible under microscope. Individual colonies were then selected and expanded in separate wells. The expression of the recombinant protein was measured by ELISA using anti‐His tag monoclonal antibody (Invitrogen) to capture and biotinylated anti‐ANG1 antibody for detection (R&D System). The clones showing the highest titers of the recombinant protein were selected. Protein production was in a Nunc CellFactory system (ThermoFisher) using serum‐free culture. The medium was centrifuged to clear the cell debris, followed by concentrating and buffer exchange into IMAC binding buffer using Vivaflow 100 system (Sartorius). The resulting protein solution was purified by Histrap column (GE Healthcare) and then buffer exchanged into DPBS by using desalting column (Thermo). For further scale‐up production of the C4BP‐ANG1 variants, an alternative CHO expression system was used, as described previously (36, 37).

### Biochemical characterization of C4BP‐ANG1

The molecular size of the protein complex was determined by gel electrophoresis, western blot and size‐exclusion chromatography. Under either reduced or non‐reduced conditions, the molecular size of the monomeric or the heptameric form was determined by SDS‐PAGE and western blot, respectively. Anti‐His tag monoclonal antibody (Invitrogen) was used in western blot analysis.

TEM analysis of the protein complexes was conducted following a standard negative staining protocol (38). In brief, purified C4BP‐A1 was diluted in DPBS to a concentration of 100 ug/mL. A 10 uL droplet was applied to a glow‐discharged carbon‐coated copper grids and allowed to sit for 1 minutes. The grids were washed by dipping in two separate drops of water followed by two drops of 2% uranyl acetate (Electron Microscopy Sciences). Grids were examined at the Northwestern Electron Probe Instrumentation Center (EPIC) using Hitachi HT‐7700 Biological S/TEM Microscope.

C4BP‐ANG1 and Tie2 ectodomain (amino acid 1‐745 of Tie2 in Tie2‐Fc fusion) binding was measured by coimmunoprecipitation assay. 100 ng recombinant Ang1‐C4BP, C4BP‐Ang1 or native Angpt1 protein (R&D systems) was mixed with 100 ng Tie2‐Fc (with His tag, R&D systems) in TBST buffer at 4°C for 4 hours. Then 10 ul protein G agarose beads (Sigma) were added and incubated for another 1 hour. The beads were collected by centrifugation and washed five times by TBST. The immunoprecipitated proteins were eluted by reducing SDS‐PAGE sample buffer and analyzed by western blot using anti‐His tag antibody.

### Cell‐based experimental studies of C4BP‐ANG1

Two experimental systems were used to evaluate cell signaling response to C4BP‐ANG1 treatment: human umbilical vein cells (HUVECs, ATCC) and HEK293 with stable expression of full length Tie2 with a C‐terminus FLAG tag. Dose and time response of HUVEC cells were conducted by adding purified C4BP‐ANG1 proteins directly to the culture medium for indicated period of time, followed by cell lysis for immunoblotting of phospho‐Akt. The HEK293 cell study was conducted with the stimulation of the cells with C4BP‐ANG1 followed by anti‐FLAG immunoprecipitation (IP) of Tie2‐FLAG that were stably expressed at the cell surface. Tie2‐FLAG from IP was then subjected to immunoblotting analysis of phospho‐Tie2 measured by anti‐phosphotyrosine antibody blotting (4G10 Platinum, Millipore).

Immunofluorescence staining of HUVEC in response to C4BP‐ANG1 was conducted as described previously (39). In brief, HUVECs were transfected by electroporation with full‐length Tie2 fused with a C‐terminus FLAG tag. Two days after transfection, purified C4BP‐ANG1 protein or buffer control was added to the culture medium. Following an incubation period of 15 min the cells were fix and then stained with anti‐FLAG antibody (Sigma) to detect Tie2‐FLAG. To investigate the integrity of cell junctions, HUVECs were treated with LPS (100 ng/mL) or VEGF (100 ng/mL) for 30 min, in presence of vehicle or C4BP‐A1 (500 ng/mL). The cells were fixed and stained with anti‐VE‐Cadherin antibody (Millipore).

### *In vivo* Tie2 activation assay

Vehicle or C4BP‐Ang1 were intravenously injected to 8 weeks old female Balb/c mice or intraperitoneally injected to P6 Balb/c neonates. After the specified time, the mice were sacrificed, and their lungs were collected and lysed by RIPA buffer supplemented with proteinase/phosphatase inhibitor cocktail (50 mg tissue per mL lysis buffer). Tie2 protein was immunoprecipitated from the lysates by using anti‐Tie2 antibody (Millipore) and protein A/G‐sepharose. The phosphorylation level of precipitated Tie2 was assessed by anti‐phosphotyrosine antibody as described above.

### Rabbit eye pharmacokinetic study

New Zealand White (NZW) rabbits were used to assess the intraocular pharmacokinetics of C4BP‐ANG1. The procedures were described previously (40). Briefly, proparacaine hydrochloride (Alcain, Alcon) was administered into both eyes to minimize discomfort. Approximately 1μg of C4BP‐ANG1 protein in 100μL DPBS solution was injected intravitreously to one of the eyes using a 700 series Hamilton syringe and a 30 gauge needle. The contralateral eye was injected with the buffer as a control. Aqueous humor was collected daily (preinjection sample: day 0) for seven consecutive days. The levels of C4BP‐ANG1 were measured by ELISA as described above. On the seventh day the animals were sacrificed and vitreous humor samples from both eyes were collected for detection of C4BP‐ANG1 levels by ELISA.

### *In vivo* assessment of C4BP‐ANG1

Miles assay for VEGF induced skin vascular leakage was performed as described previously (41). Briefly, 8 weeks old female Balb/c mice were *i.p.* injected with histamine inhibitor pyrilamine (40 ul/g body weight) followed by tail vein injection of 100 μL Evans blue dye (1% in PBS). After 30 min, the mice were anesthetized by Ketamine/Xylazine cocktail. Then vehicle or VEGF (100ng, R&D Systems), either in the presence or the absence of C4BP‐Ang1 (250 ng), was intradermally injected into different sites of the mouse flank. The mice were sacrificed 20 min after the intradermal injections and the leakage of Evans blue was photographed and then quantified by deionized formamide extraction followed by spectrophotometer measurement on 620 nm.

To investigate whether pretreatment with C4BP‐Ang1 prevents VEGF induced skin vascular leakage, two groups of 10 weeks old female Balb/c mice received tail vein injection of either Evans blue or Evans blue solution with C4BP‐Ang1 (0.5 ug/g body weight). After 30 min, VEGF at different doses were intradermally injected into different sites of the mouse flank. The mice were sacrificed 20 min after the intradermal injections and the leaked Evans blue in each site was photographed.

In the mustard oil induced vascular leakage model, 8 weeks old female Balb/c mice were *i.v.* injected with either vehicle or C4BP‐ANG1 protein. After 30 min, all the mice received *i.v.* injection of 100 μL Evans blue dye (1% in DPBS). 10 uL Mustard oil (5% allyl‐isoyhiocyanate diluted in mineral oil; Sigma) or mineral oil control was then applied to the dorsal and ventral surfaces of the ears. After 20 min, the ears were collected and the leaked Evans blue was quantified as described above.

In LPS‐induced lung vascular leakage model, 8 weeks old female Balb/c were anesthetized by Ketamine/Xylazine cocktail and randomly divided into 3 groups: (1) The control group: Mice received intranasal inhalation of 20 uL PBS and *i.v.* injection of PBS. (2) LPS group: Mice received intranasal inhalation of 20 uL LPS (1 ug/uL, from Escherichia coli 0111:B4, Sigma) and i.v. injection of PBS. (3) LPS + C4BP‐A1 group: Mice received intranasal inhalation of 20 uL LPS and *i.v.* injection of C4BP‐A1 (0.5 ug /g BW, in PBS). Lung permeability was assessed by *i.v.* injecting of 100 uL Evans blue (1% in PBS). One hour after Evans blue administration, the mice were euthanized and the lung tissues were collected. The leakage of Evans blue was quantified as described above.

## Acknowledgments

This study was funded by National Institutes of Health (R01EY025799 to S.E.Q.) and by Mannin Research, Inc. (to S.E.Q and J.J.). We are grateful to Dr. Yves Durocher and his team at the National Research Council Canada for recombinant protein production and analysis. We thank Emily Anne Hicks for technical assistance to the rabbit eye study, and George Nikopoulos for his suggestions and for facilitating the collaborations.

