## Supplemental Figures for "New Soluble Angiopoietin Analog of C4BP-ANG1 Prevents Pathological Vascular Leakage"

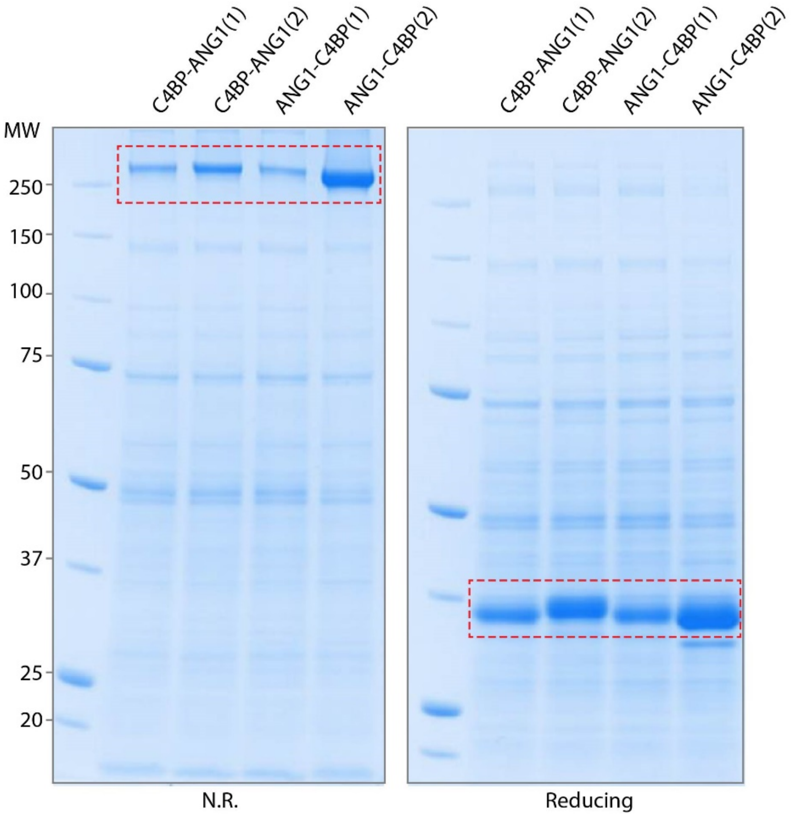


**Supplementary Figure S1. C4BP and ANG1 fusions in various construction formats all form heptamers in near homogeneity.** In an N-to-C-terminus order, 4 plasmids for mammalian cell expression were constructed: 1. C4BP-ANG1(1) with a C-terminus 6xHis tag, 2. C4BP-ANG(2) with an N-terminus 6xHis tag, 3. ANG1-C4BP(1) with a C-terminus 6xHis tag, and 4. ANG1-C4BP(2) with an N-terminus 6xHis tag. Proteins were expressed in CHO cells cultured in serum-free medium and subsequently harvested from the culture medium. By performing SDS-PAGE analysis of the media under either non-reducing (N.R.: left panel) or reducing (right panel) conditions, it was determined that C4BP-ANG1 proteins (highlighed in red boxes), regardless of their N-to-C orders, naturally form heptamers (with multiplicity of 7) of ~280 kDa via disulfide bridges. All fusion proteins can be reduced to their ~35 kDa monomeric forms under reducing condition.

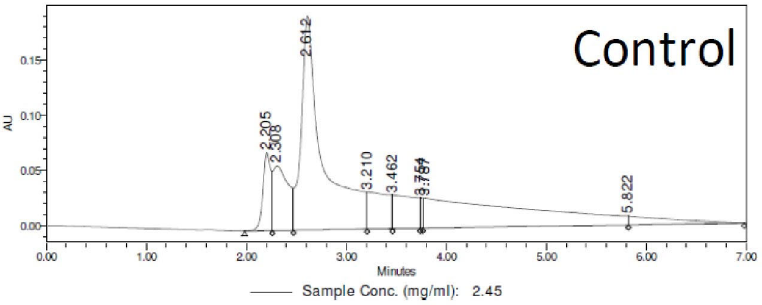


| | RT | % Area | Area | Height |
|---|---|---|---|---|
| 1 | 2.205 | 5.36 | 388192 | 70372 |
| 2 | 2.308 | 8.75 | 633834 | 58511 |
| 3 | 2.612 | 40.38 | 2924070 | 194026 |
| 4 | 3.210 | 6.70 | 485109 | 33664 |
| 5 | 3.462 | 6.63 | 480258 | 30660 |
| 6 | 3.754 | 0.63 | 45418 | 27481 |
| 7 | 3.787 | 27.96 | 2024484 | 27126 |
| 8 | 5.822 | 3.58 | 259248 | 7986 |

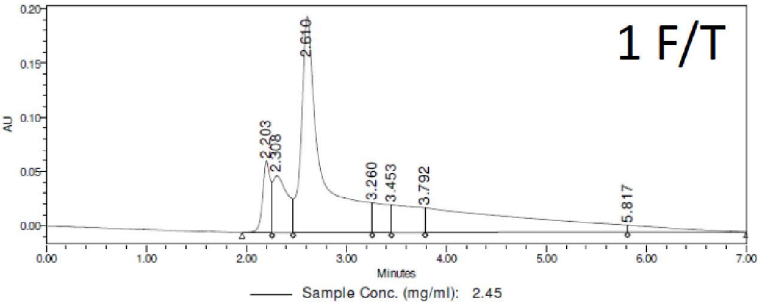


| | RT | % Area | Area | Height |
|---|---|---|---|---|
| 1 | 2.203 | 5.67 | 361276 | 66147 |
| 2 | 2.308 | 8.53 | 543510 | 52450 |
| 3 | 2.610 | 43.94 | 2801454 | 199371 |
| 4 | 3.260 | 4.74 | 302266 | 27484 |
| 5 | 3.453 | 7.67 | 488655 | 25646 |
| 6 | 3.792 | 26.06 | 1661291 | 22888 |
| 7 | 5.817 | 3.40 | 216640 | 6724 |

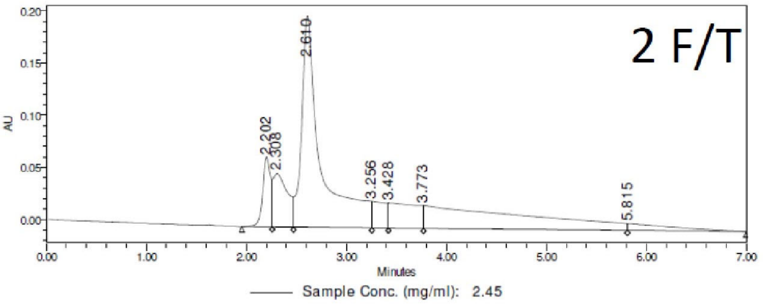


| | RT | % Area | Area | Height |
|---|---|---|---|---|
| 1 | 2.202 | 5.92 | 363568 | 66921 |
| 2 | 2.308 | 8.50 | 521839 | 50984 |
| 3 | 2.610 | 44.10 | 2708890 | 202368 |
| 4 | 3.256 | 3.96 | 243471 | 25479 |
| 5 | 3.428 | 7.75 | 475900 | 23924 |
| 6 | 3.773 | 26.46 | 1625103 | 21823 |
| 7 | 5.815 | 3.32 | 203823 | 6596 |

**Supplementary Figure S2. Protein stability following freeze-thaw cycles.** Purified ANG1-C4BP(1) was subjected to one or two freeze-thaw cycles (F/T) before UPLC-SEC analysis of heptamer quality (at peak 2.610). No loss of the heptamer fraction was apparent (compare 1 F/T and 2 F/T with the control that was stored at 4 °C).
